# Structural variants are a major source of gene expression differences in humans and often affect multiple nearby genes

**DOI:** 10.1101/2021.03.06.434233

**Authors:** Alexandra J. Scott, Colby Chiang, Ira M. Hall

## Abstract

Structural variants (SVs) are an important source of human genome diversity but their functional effects are not well understood. We mapped 61,668 SVs in 613 individuals with deep genome sequencing data from the GTEx project and measured their effects on gene expression. We estimate that common SVs are causal at 2.66% of eQTLs, which is a 10.5-fold enrichment relative to their abundance in the genome and consistent with prior work using smaller sample sizes. Duplications and deletions were the most impactful variant types, whereas the contribution of mobile element insertions was surprisingly small (0.12% of eQTLs, 1.9-fold enriched). Multi-tissue analysis of expression effects revealed that gene-altering SVs show significantly more constitutive effects than other variant types, with 62.09% of coding SV-eQTLs active in all tissues with known eQTL activity compared to 23.08% of coding SNV- and indel-eQTLs, whereas noncoding SVs, SNVs and indels show broadly similar patterns. We also identified 539 rare SVs associated with nearby gene expression outliers. Of these, 62.34% are noncoding SVs that show strong effects on gene expression yet modest enrichment at known regulatory elements, demonstrating that rare noncoding SVs are a major source of gene expression differences but remain difficult to predict from current annotations. Remarkably, both common and rare noncoding SVs often show strong regional effects on the expression of multiple genes: SV-eQTLs affect an average of 1.82 nearby genes compared to 1.09 genes affected by SNV- and indel-eQTLs, and 21.34% of rare expression-altering SVs show strong effects on 2-9 different genes. We also observe significant effects on gene expression extending 1 Mb from the SV. This provides a mechanism by which individual noncoding SVs may have strong and/or pleiotropic effects on phenotypic variation and disease.

## INTRODUCTION

Structural variants (SVs) are a diverse class of genetic variation that include copy number variants (CNVs), mobile element insertions (MEIs) and balanced rearrangements at least 50 base pairs (bp) in length. While SVs are relatively rare compared to single-nucleotide variants (SNVs) and small insertion or deletion (indel) variants, their size and diversity mean that SVs can disrupt protein-coding genes and genomic regulatory elements through diverse mechanisms. Furthermore, SVs often have more severe consequences compared to smaller variants and previous studies have found that SVs have an outsized impact on human gene expression compared to their relative abundance in the genome (Chiang et al. 2017; Stranger et al. 2007; Sudmant et al. 2015). SVs have also been implicated in the biology of human diseases such as autism spectrum disorder (Brandler et al. 2018; Sebat et al. 2007; Turner et al. 2017; Weiss et al. 2008) and schizophrenia (International Schizophrenia Consortium 2008; Marshall et al. 2017; McCarthy et al. 2009; Walsh et al. 2008). However, SVs are difficult to detect from short-read DNA sequencing data and are often excluded from complex trait association studies.

Advances in high-throughput sequencing technologies that have allowed for widespread use of whole genome sequencing (WGS), combined with advances in scaling SV detection algorithms, mean that comprehensive studies of all forms of genetic variation are now possible for large human cohorts. Recent studies of SV in large, deeply-sequenced human cohorts have found that SVs account for 4.0-11.2% of rare high-impact coding alleles (Abel et al. 2020) and are responsible for 25-29% of rare protein-truncating events per genome (Collins et al. 2020). However, few studies to date have examined the functional effects of SV on gene expression and these studies are limited to relatively small cohort sizes or only a few tissue types with available gene expression data (Chiang et al. 2017; Han et al. 2020; Jakubosky et al. 2020; Sudmant et al. 2015).

Here, we use deep WGS data and multi-tissue RNA-seq expression data from 613 individuals in the Genotype-Tissue Expression (GTEx) project to comprehensively map SVs and to evaluate their impact on both common and rare gene expression changes across 52 tissue types. This study expands on our prior analysis of SV in 147 human samples from the GTEx cohort with RNA-seq expression data from 13 different tissues (Chiang et al. 2017) and is the most comprehensive study of SV-eQTLs to date. The expanded cohort size provides greater power to evaluate the impact and mechanisms of SV-associated gene expression changes, particularly for rare SVs. We found that both rare and common SVs have broad effects, often affecting more genes and having larger effect sizes compared to SNVs and indels, and that these effects are frequently caused by noncoding SVs.

## RESULTS

### Variant calling

We mapped structural variation (SV) in 613 individuals from the GTEx v7 release using LUMPY (Chiang et al. 2015; Layer et al. 2014), svtools (Larson et al. 2019), GenomeSTRiP (Handsaker et al. 2011, 2015) and the Mobile Element Locator Tool (MELT) (Gardner et al. 2017) (see **Methods**). Variant calls were filtered and merged using the same approach as in our previous GTEx study (Chiang et al. 2017; Li et al. 2017), resulting in a total of 61,668 “high confidence” SVs that are the basis for all subsequent analyses (**Table 1**). Single nucleotide (SNV) and small insertion deletion (indel) variants were mapped using GATK (McKenna et al. 2010) as part of the official v7 release from the GTEx Consortium.

**Table 1.** Summary of variant types and eQTL mapping. SVs were detected based on breakpoint evidence (BP) or read-depth evidence (RD). SNVs and indels were called using the Genome Analysis Toolkit (GATK). Common variants (MAF ≥ 0.01) were used to map cis-eQTLs.

|  | Detection method | No. variants | Median size (bp) | # of common variants | eVariants |
| --- | --- | --- | --- | --- | --- |
| SNV | GATK | 37,087,030 | 1 | 9,609,545 | 178,000 |
| Indel | GATK | 3,081,270 | 3 | 818,401 | 16,460 |
| Deletion (DEL) | BP | 20,954 | 1,311 | 4,385 | 210 |
|  | RD | 10,252 | 2,151 | 8,166 | 66 |
| Duplication (DUP) | BP | 3,388 | 2,632 | 1,090 | 64 |
|  | RD | 1,598 | 6,891 | 896 | 233 |
| Multi-allelic CNV (mCNV) | RD | 4,365 | 3,602 | 3,238 | 460 |
| Inversion | BP | 295 | 1,054 | 96 | 2 |
| Reference mobile element insertion (MEI-del) | BP | 2,681 | 306 | 2,026 | 88 |
| Non-reference mobile element insertion (MEI-ins) | BP | 13,066 | 280 | 4,496 | 91 |
| Other (BND) | BP | 5,069 | - | 2,010 | 57 |
| All SVs | - | 61,668 | - | 26,409 | 1,271 |
| All Variants | - | 40,229,968 | - | 10,454,355 | 195,731 |

### Effects of common SVs

We performed *cis*-eQTL mapping of common variants (MAF ≥ 0.01) using a permutation-based mapping approach with FastQTL (Ongen et al. 2016), limiting comparisons to variants within 1 Mb of the transcription start site (TSS) of each gene. We performed eQTL analyses in each of the 48 tissues for which expression data was available for at least 70 individuals and defined an eQTL as an eVariant/eGene pair detected in a given tissue. We performed a “joint” eQTL mapping analysis in which SVs, SNVs and indels were simultaneously queried for eQTL status, allowing for direct comparisons between their properties and identification of a likely causal variant. An SV was the lead marker in 2.66% (7,960/299,187) of eQTLs (**Supplemental Table S1**), although this is likely an underestimate of SV causality due to inferior genotyping accuracy for SVs, which biases eQTL fine-mapping analyses against SVs. While this estimate of the contribution of SVs is relatively small, it represents an 10.5-fold enrichment over the abundance of SVs in the genome. This result is consistent with our prior analysis of the initial 147 individuals from the GTEx cohort (Chiang et al. 2017).

A novel aspect of this study is that we used MELT to sensitively map mobile element insertion (MEI) variants, including non-reference insertions that were not detected in our prior GTEx studies. It has been proposed that MEIs may have broad effects on gene expression due to their ability to disrupt genes, promote epigenetic gene silencing, and serve as alternate promoters (Payer and Burns 2019; Chuong et al. 2017); however, there has been scant data in humans to address this. We found that only 0.12% (353/299,187) of eQTLs had an MEI as the lead marker. Although this is still a 1.9-fold enrichment of predicted causal MEIs relative to their abundance (0.06% of common variants), MEIs were far less likely than other SV types to be the lead marker (e.g., mCNVs are enriched 45-fold, duplications 38-fold and deletions 3.3-fold). Thus, despite compelling molecular evidence for the functional potential of MEIs, our results suggest that they are only slightly enriched as causal eQTL variants relative to SNVs and indels and are depleted relative to other SVs, on average.

Interestingly, we found that not only do SVs have larger effect sizes compared to SNPs and indels, as noted in previous studies (**Supplemental Fig. S1**) (Jakubosky et al. 2020; Chiang et al. 2017), they are also more likely to alter the expression of multiple nearby genes. Each eSV affects an average of 1.82 unique eGenes while SNVs and indels affect an average of 1.09 unique eGenes. Although this effect is partially explained by large SVs that alter the copy number of multiple adjacent genes, there is also a significant difference for genes affected by noncoding eVariants: on average, eSVs affect 1.50 unique eGenes for which they do not intersect any exons of the eGene, compared to an average of 1.04 unique eGenes for SNVs and indels (p=1.02×10^−55^, one-sided Mann-Whitney U test) (**1B-D)**. These noncoding effects are most pronounced for duplications (p=6.10×10^−53^) and mCNVs (p=4.75×10^−56^), which are the only two categories of noncoding SVs that affect significantly more eGenes than point variants. This result indicates that causal SVs are generally more impactful than causal point variants, both in terms of their per-gene effect sizes as well as their potential to affect multiple genes. These results also suggest that SVs are more likely to disrupt key regulatory elements and/or alter higher-order genome architecture, allowing individual variants to affect multiple genes.

To investigate the functional mechanisms of expression-altering SVs, we defined a set of putative causal SVs using a score generated by taking the product of the causal probability calculated using CAVIAR (Hormozdiari et al. 2014) and the fraction of heritability attributed to the SV calculated using GCTA (Yang et al. 2011) (**Supplemental Table S2**), as described previously (Chiang et al. 2017). At each eGene we selected the SV within the *cis*-region that had the strongest association with the eGene’s expression and allocated these 10,911 unique SVs into six bins on the basis of causality score quantiles, with the least-causal bin containing the 50% of SVs with the lowest scores. Next, we measured the enrichment of SVs in each causality bin at a diverse set of genomic annotations and in the core 15 chromatin segmentation states from the Roadmap Epigenomics Project using a permutation test based on shuffled genomic positions (**Supplemental Fig. S2-S3; see Methods)**. SVs in the most causal quantiles were strongly enriched in the exons of their associated eGenes, which is expected and confirms that our causality score is informative. We also observed an enrichment of causal SVs in the 10 kb regions upstream of the TSS and downstream of the 3’ UTR. Additionally there is a small enrichment of the causal SVs in segmental duplications, which is likely driven by large mCNVs at multi-copy genes. However, predicted causal SVs were not enriched in any other genomic features tested, which suggests that while eSVs are generally found relatively close to their eGenes, they may be altering expression through diverse mechanisms and that our study is underpowered to identify enrichments in specific regulatory element classes. Alternatively, existing annotations may be insufficiently informative to detect functional enrichments for the variants and tissues analyzed here.

The number and diversity of tissues with expression data allows us to evaluate the tissue specificity of eQTLs. We hypothesized that SVs might have more ubiquitous effects on gene expression than point variants due to constitutively-acting dosage changes or due to complete deletion or duplication of regulatory elements rather than more subtle effects, for example, on transcription factor binding. To allow for facile comparisons between variant types, we limited this analysis to variant-gene pairs with a significant association in our eQTL analysis for which expression data was available across all 48 tissues. We used METASOFT (Han and Eskin 2011) to evaluate eQTL activity across all 48 tissues and evaluated eQTLs for which active (*m*<0.9) or inactive (*m*<0.1) status could be determined in at least 43 tissues. We found that coding SV-eQTLs are more constitutive than other eQTL classes, showing activity across a larger proportion of tissues compared to SNV- and indel-eQTLs (**Fig. 1E**). Whereas 92.16% of coding SV-eQTLs are constitutively active – defined here as active in >75% of tissues with known status – only 74.12% of coding SNV- and indel-QTLs are constitutive. However, the result at noncoding eQTLs is less clear: while 44.44% of noncoding SV-eQTLs are active in 100% of tissues with known activity compared to 26.23% of noncoding SNV- and indel-eQTLs, the difference does not hold if we examine eQTLs that are active in >75% of tissues with known status (74.86% of noncoding SV-eQTLs are constitutive; 74.12% of noncoding SNV- and indel-eQTLs are constitutive) (**Supplemental Fig. S4**). This shows that coding SVs typically impact expression across many tissues, whereas smaller and noncoding variants tend to affect gene expression on a more tissue-specific basis. In contrast to coding SV-eQTLs, noncoding SV-eQTLs show similar patterns of tissue specificity to noncoding SNV- and indel-eQTLs, indicating that these variant types are likely to function through similar mechanisms. However, it is important to note that noncoding SV-eQTL activity could not be determined by METASOFT in many tissues (**Supplemental Fig. S5**), so it is possible that the true tissue specificity of noncoding SVs may differ from noncoding SNVs and indels. This appears to be the result of relatively large effect-size standard errors for SV-eQTLs that result from genotyping inaccuracies. While METASOFT can determine cross-tissue eQTL activity when effect sizes are large despite large standard errors, as seen in coding SV-eQTLs, when effect sizes are small but effect size errors are large, the algorithm often cannot confidently judge activity (**Supplemental Fig. S6**).

**Figure 1.**
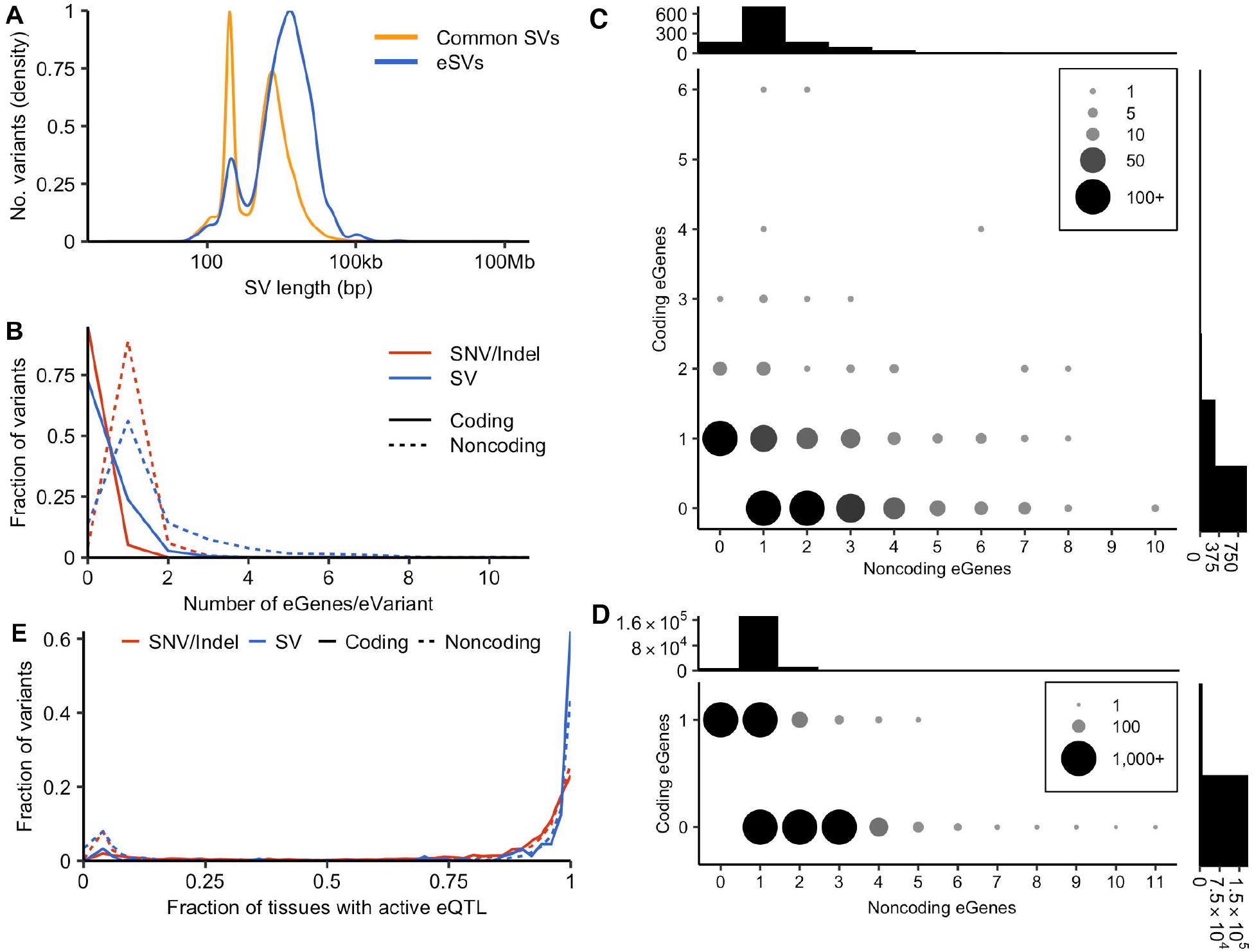
Features of SV-eQTLs. (**A**) Size distribution of eSVs (blue line) compared to all common SVs (yellow line). (**B**) Distribution of the number of eGenes per eVariant for SVs (blue lines) compared to SNVs and indels (red lines). Solid lines indicate the number of “coding” eGenes whose coding regions are intersected by the associated eVariant and dashed lines indicate the number of “noncoding” eGenes that are not intersected by the associated eVaraint. (**C,D**) The number of eVariants with the indicated combination of coding and noncoding eGenes, as defined above. Shown for SVs (**C**) and SNV/indels (**D**), with histograms showing the total number of eVariants with the indicated number of associated coding or noncoding eGenes above the y- and x-axes, respectively. (**E**) Distribution of the tissue specificity of eQTLs across tissues, as evaluated by METASOFT, for eQTLs in which the activity status is known in at least 43 of 48 evaluated tissues. Red lines indicate the distribution of SV-eQTLs that are active in the fraction of evaluated tissues indicated on the x-axis. Blue lines indicate the same for SNV- and indel-eQTLs. Solid lines denote coding eQTLs where the eVariant intersects the coding region of the associated eGene and dashed lines show the distributions for noncoding eQTLs.

### Effects of rare SVs

Rare SVs are enriched near genes with highly aberrant expression (Chiang et al. 2017) and are more likely to have large effect sizes compared to other variant types (Li et al. 2017). To assess the effects of rare SVs on gene expression, we identified genes in which individuals displayed highly aberrant gene expression levels compared to the dataset as a whole. We limited this analysis to the 513 individuals of European descent to reduce the effects of population stratification. We defined 27,292 multi-tissue gene expression outliers (median |Z| ≥ 2 across all tissues in an individual) and 346,122 “tissue-restricted” outliers with highly aberrant expression (|Z|≥4) in two or more tissues in the same individual. Next, we identified 13,768 “singleton” SVs smaller than 1 Mb in size that were positively genotyped in one individual. These rare SVs are strongly enriched within the gene body and flanking sequence of multi-tissue gene expression outliers compared to the null expectation in 1,000 random permutations of the outlier sample names, with enrichment decreasing as flanking distance increases (**Supplemental Fig. S7**). The enrichment of rare SVs in close proximity (i.e., within 5 kb) to multi-tissue gene expression outliers is consistent with our prior work (Chiang et al. 2017), but the increased power in this study allows us to observe enrichment at greater distances as well. At flanking distances as large as 50 kb, we observe a 6.4-fold enrichment of rare SVs around multi-tissue outliers, suggesting that rare SVs contribute to rare expression differences even from relatively large genomic distances. Importantly, because gene expression values can only decrease to 0, a conservative Z-score limit such as the one used for tissue-restricted outliers favors gene expression outliers with increased expression, thus limiting our ability to detect SVs associated with decreased expression (**Supplemental Fig. S8**). However, these conservative outlier definitions, combined with the above enrichment results, provide confidence in the set of outlier-associated SVs.

A total of 539 unique outlier-associated SVs are located in the gene body and 50 kb flanking region of gene expression outliers (**Fig. 2A**; **Supplemental Table S3**). Notably, 62.34% (336/539) of these are noncoding SVs that do not affect the coding sequence of one or more expression outliers, and 0.80% of expression outliers are associated with noncoding SVs. This contradicts the general assumption that rare SVs typically act through gene dosage effects. Duplications and mCNVs are most likely to be associated with an expression outlier, MEIs are least likely (**Fig. 2B**) and larger SVs are more likely to be outlier-associated regardless of type (**Fig. 2C**). However, many outlier-associated SVs are smaller in size (**Fig. 2D**). For example, 13.33% (50/375) of SVs associated with tissue-restricted outliers are smaller than 1 kb and nearly half (49.33%; 185/375) are smaller than 10 kb. Multi-tissue outlier-associated SVs tend to be slightly larger, with only 4.98% (12/241) smaller than 1 kb and 35.27% (85/241) smaller than 10 kb. Overall these results provide further evidence that rare SVs often affect gene expression through more complex mechanisms than large, dosage-altering events.

**Figure 2.**
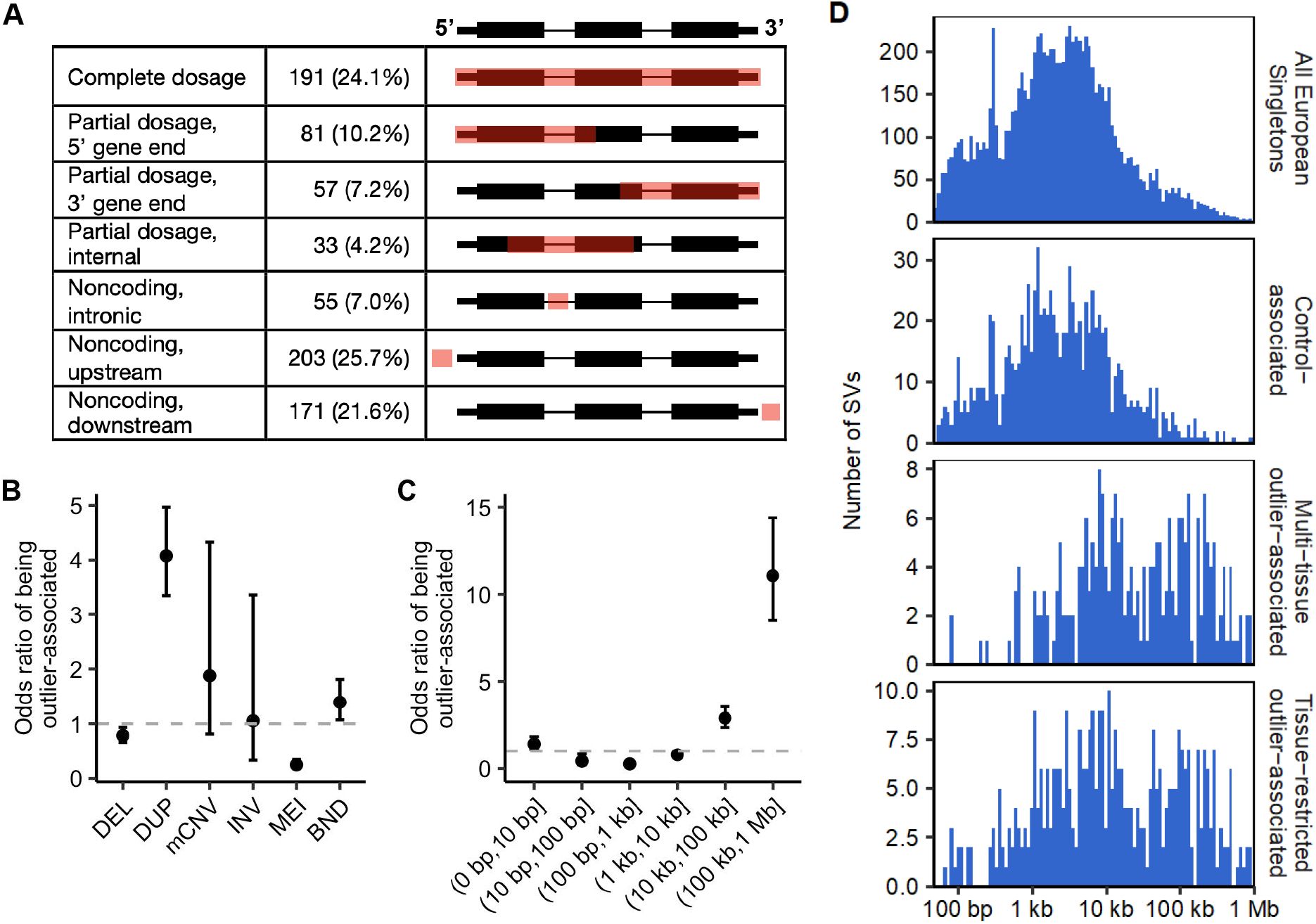
Features of outlier-associated SVs. (**A**) Location of outlier-associated SVs relative to their associated outlier gene and the number of SV/outlier gene associations identified in each category. Percentages indicate the fraction of outlier/SV pairs found at each relative location compared to the total number of SV/outlier gene associations. Note that this definition means that one SV may be associated with multiple outlier genes and thus is counted in multiple categories. Gene diagrams provide examples of possible SV location, shown in red, relative to the outlier gene. (**B,C**) Odds ratio of being outlier-associated by SV type (**B**) and SV size (**C**). (**D**) Distribution of SV sizes for singleton SVs smaller than 1 Mb identified in European individuals that were used in outlier analyses. Panels depict size distributions for all European-cohort singletons, control-associated singletons, multi-tissue outlier-associated singletons and tissue-restricted outlier-associated singletons.

We next sought to determine if rare outlier-associated SVs are enriched in annotated genomic features. Although there was little signal in our enrichment analysis of common SVs, as described above, rare variants typically have larger effect sizes and are more likely to be deleterious. For this analysis, we defined a set of “control” SVs that are located within or near genes but do not exhibit expression effects. We identified 1,405 singleton SVs (1,327 noncoding) located within 50 kb of genes that showed consistent expression levels (|Z| < 1) across all tissues in an individual. Although this is not an ideal set of control SVs considering that some may in fact alter gene expression in tissues or development timepoints for which expression was not measured, it is nonetheless a relatively conservative set of likely-nonfunctional SVs that can be used for comparison to outlier-associated SVs. We examined the overlap of both outlier- and control-associated noncoding SVs with annotated genomic features and with segmentation states from the Roadmap Epigenomics Project core 15-state model (**Fig. 3A**). We observed significant enrichment of outlier-associated SVs in 5 of the 34 evaluated features and chromatin states (Fisher’s Exact test; Bonferonni p < 0.05). Most of these significant associations are in Roadmap Epigenomics Project segmentation states in close proximity to transcribed genes, including transcription at the 5’ and 3’ end of genes showing both promoter and enhancer signatures, active transcription start sites and regions flanking active transcription start sites. We also observed significant enrichment in the Roadmap Epigenomics Project segmentation state associated with zinc finger protein genes and in enhancer annotations from Genehancer. It is important to note, however, that the number of overlaps observed in this analysis is small and increased power might change these results. Thus, while rare SVs appear to have dramatic effects on gene expression, most existing functional annotations are not very informative. Consistent with this, the distribution of SV impact scores (Ganel et al. 2017) is not significantly different between expression-altering SVs and control SVs (**Supplemental Fig. S9**).

**Figure 3.**
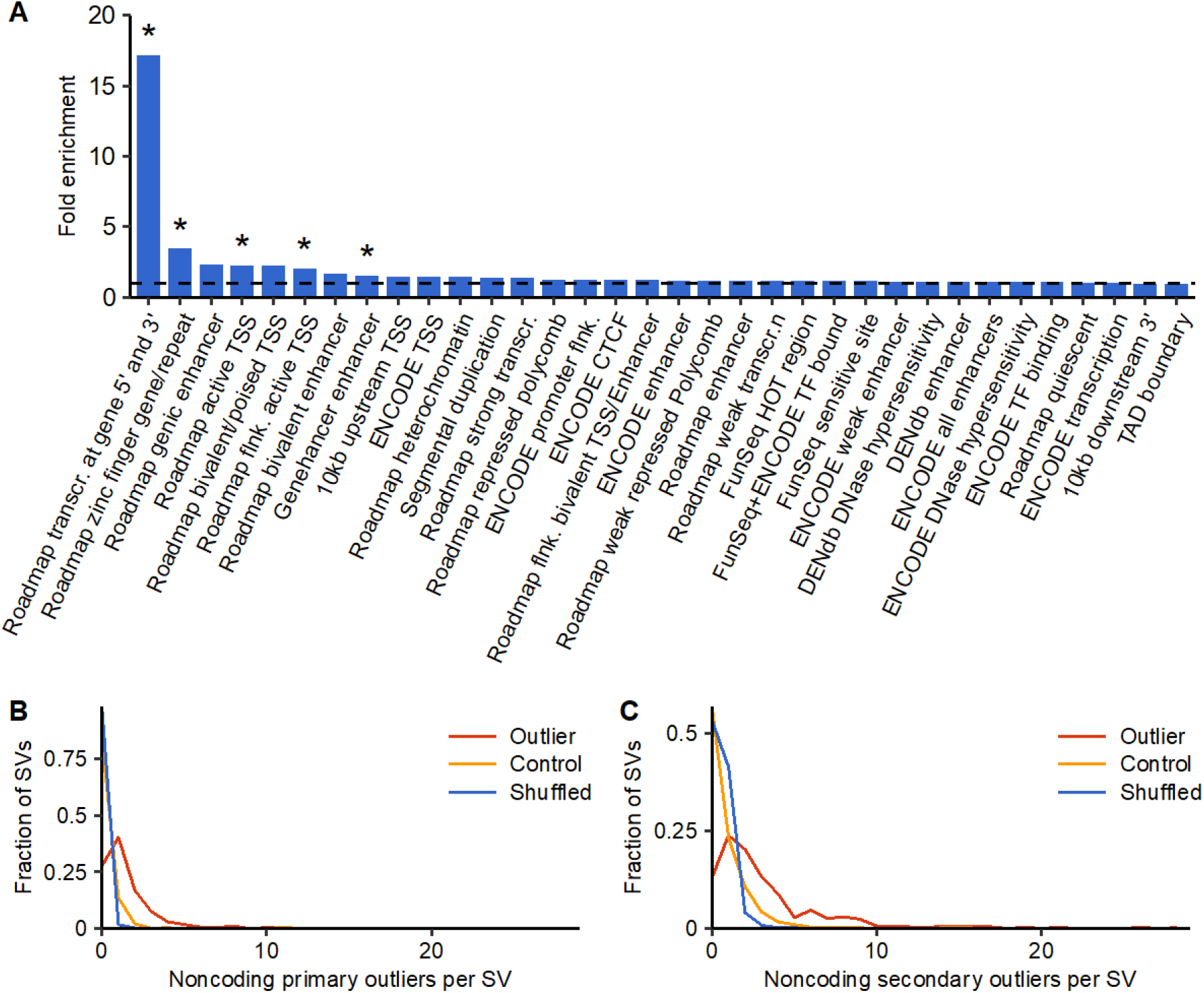
Mechanistic insights into outlier-associated SVs. (**A**) Enrichment of outlier-associated SVs in functional genomic annotations compared to control-associated SVs. Asterisks indicate statistical significance based on a Fisher’s exact test with Bonferroni correction for multiple testing. (**B,C**) The distribution of the number of noncoding primary (**B**) and secondary (**C**) outliers found within 1 Mb of the region surrounding tissue-restricted outlier-associated SVs (red), control-associated SVs (yellow) and a shuffled null (blue).

Interestingly, we found that 115 (21.34%) of outlier-associated SVs are associated with more than one expression outlier, and that 8 are associated with 5-9 expression outliers, suggesting that many rare SVs may have regional effects. In order to evaluate these broader regional effects of rare expression-altering SVs, we relaxed the definition for aberrant expression to generate a set of “secondary” expression outliers in which the tissue-restricted (“primary”) outlier absolute Z-score cutoff was reduced to 3 in at least two tissues. We found significantly more primary and secondary outliers within 1 Mb of the 469 tissue-restricted outlier-associated SVs compared to the 1,496 control-associated SVs and to a shuffled null in which we randomly shuffled the sample names of outlier-associated SVs 1,000 times and counted the median number of associated outlier genes (**3B,C**). This increase is especially pronounced for secondary outliers whose coding regions do not overlap with the associated SV. We observe that noncoding outlier-associated SVs are associated with an average of 1.44 primary outliers (|Z|≥4) compared to an average of 0.02 associated primary outliers surrounding the shuffled null SVs (p-value=2.78×10^−106^; one-sided Mann-Whitney U test). These differences remain for secondary outliers, with an average of 3.34 secondary outliers found in the expanded region surrounding noncoding outlier-associated SVs compared to an average of 0.54 secondary outliers for the shuffled null (p-value=4.94×10^−76^; one-sided Mann-Whitney U test). These results suggest that rare SVs have far-reaching effects on gene expression and that these effects are primarily driven by noncoding regulatory mechanisms rather than changes to gene copy number.

## DISCUSSION

We have comprehensively mapped SVs from WGS data in 613 individuals from the GTEx dataset and analyzed the impact of both common and rare SV on human gene expression across 53 tissue types. Our findings confirm results from previous analyses that SVs make an outsized contribution to common gene expression changes compared to their abundance in the genome and play an important role in rare gene expression differences (Chiang et al. 2017). A novel aspect of this study is the inclusion of a comprehensive set of MEI insertions, including those present in the GTEx samples but not the reference genome. We observed that MEIs do not play an especially important role in determining gene expression differences. In contrast, we found that mCNVs play an extremely impactful role, being 45-fold enriched among eQTL lead markers compared to their abundance in the genome and more likely to be associated with gene expression outliers (OR=1.88). mCNVs were found to give rise to most human variation in gene dosage (Handsaker et al. 2015), but our findings indicate that noncoding functional mCNVs are also abundant in the human genome.

One of the major motivators for studies such as this one is to understand the role of genetic variation in affecting gene transcription. Unfortunately, expression-altering SVs were not well correlated with any specific functional annotations other than proximity to genes, and thus existing annotations are unlikely to be informative for modeling functional variant effects. This may simply be due to a lack of power given that SVs are such a diverse class of variants that can affect large genomic segments and have the potential to affect gene expression through diverse mechanisms, and our sample size is limited to 11,026 common SVs and 539 rare SVs predicted to be functional. Alternatively, the annotations currently available may be inadequate.

Nonetheless, it is clear that SVs have broad regional impacts on human gene expression, with individual variants frequently affecting multiple genes. Interestingly, these effects are not driven by large CNVs that alter the dosage of multiple coding sequences, as one might naively expect, but are most commonly observed for noncoding variants, where common noncoding eSVs affect an average of 1.50 unique genes and rare noncoding SVs associated with an average of 1.44 primary expression outlier genes. This observation suggests a mechanism by which rare noncoding SVs may be especially deleterious, and may help explain why prior work has estimated that a surprisingly large number of rare noncoding deletions – an average of 19.1 per individual – appear to be under strong purifying selection (Abel et al. 2020). Furthermore, the burden of *de novo* CNVs has been associated with autism spectrum disorder, including for noncoding variants (Turner et al. 2017; Turner and Eichler 2019). Our results provide a mechanism through which individual noncoding SVs can have strong and potentially pleiotropic effects, and thus a higher potential to contribute to disease.

While this study represents the most comprehensive analysis of the impact of SVs on human gene expression to date, our callset is missing some of the most repetitive classes of SV, such as short tandem repeats. As long read sequencing and variant calling methods improve, we will be able to gain additional insights into repetitive variants in the most complex regions of the genome. However, despite the limitations of short-read sequencing data, this study demonstrates the importance of comprehensive variant detection when evaluating genomic variants that contribute to gene expression and disease. SVs have a disproportionately large effect on common and rare gene expression changes and often affect multiple genes. Our findings reinforce the importance of comprehensive variant detection in the design of future trait mapping studies.

## METHODS

### Call set generation

We obtained 613 whole-genome BAM files from the GTEx v7 release (dbGap accession phs000424.v7.p2, accessed 1 June 2016). Structural variant calls were generated using both the SpeedSeq v0.1.1 pipeline (Chiang et al. 2015), which performs sample-level breakpoint detection via LUMPY v.0.2.13 (Layer et al. 2014) followed by population-scale merging and genotyping of SV calls via svtools v0.3.1 (Larson et al. 2019) and the GenomeSTRiP v2.00.1636 read-depth analysis pipeline (Handsaker et al. 2011), as described in our preliminary GTEx study (Chiang et al. 2017). GenomeSTRiP false discovery rate (FDR) was evaluated based on available Illumina Human Omni 5M gene expression array data (n=161) using the GenomeSTRiP IntensityRankSumAnnotator. We limited GSCNQUAL to ≥ 1 for GenomeSTRiP deletions and to ≥8 for multiallelic copy number variants, corresponding to an FDR of 10%. The GSCNQUAL cutoff for GenomeSTRiP duplications was set at ≥17, the point at which the FDR plateaued at 15.1% and did not fluctuate more than ±1% for over 50 steps of increasing GSCNQUAL score. Redundant Lumpy and GenomeSTRiP calls were merged as previously described (Chiang et al. 2017). Additionally, we ran the Mobile Element Locator Tool (MELT) v2.1.4 using MELT-SPLIT to identify ALU, SVA and LINE1 insertions into the test genomes (Gardner et al. 2017). We retained MELT calls categorized as “PASS” in the VCF info field that had an ASSESS score ≥3 and SR count ≥3. Genome Analysis Toolkit (GATK) HaplotypeCaller v3.4 (McKenna et al. 2010) SNV and indel calls were obtained from the GTEx consortium (dbGap accession phs000424.v7.p2, accessed 1 June 2016). We use allele balance instead of genotype for most analyses described in this paper because it is tolerant to alignment inefficiencies for the alternate SV allele. For MEIs identified by MELT, we converted generated genotypes (0/0, 0/1, 1/1) to integer values (0, 1, 2) that were used as a proxy for allele balance to allow for comparable analyses on these variants.

### Common eQTL mapping

We mapped *cis-*eQTLs in each of the 48 tissues for which both WGS data and RNA-seq data was available in ≥70 individuals. These tissues included: subcutaneous adipose, visceral adipose (omentum), adrenal gland, aortic artery, coronary artery, tibial artery, amygdala, anterior cingulate cortex (BA24), caudate (basal ganglia), cerebellar hemisphere, cerebellum, cortex, frontal cortex (BA9), hippocampus, hypothalamus, nucleus accumbens (basal ganglia), putamen (basal ganglia), spinal cord (cervical c-1), substantia nigra, mammary tissue, EBV-transformed lymphocytes, transformed fibroblasts, sigmoid colon, transverse colon, esophagus gastroesophageal junction, esophagus mucosa, esophagus muscularis, heart atrial appendage, heart left ventricle, liver, lung, minor salivary gland, skeletal muscle, tibial nerve, ovary, pancreas, pituitary, prostate, skin (not sun exposed), skin (sun exposed), small intestine terminal ileum, spleen, stomach, testis, thyroid, uterus, vagina and whole blood. We refer to EBV-transformed lymphocytes and transformed fibroblasts as tissue types throughout this study for convenience. Biospecimen collection, RNA-seq data alignment, RPKM calculations and data normalization were previously described (Lappalainen et al. 2013; Chiang et al. 2017).

We selected common genetic markers, defined as having MAF ≥ 0.01, for eQTL mapping. We performed a joint *cis*-eQTL analysis that included 26,409 common SVs, as well as 9,609,545 common SNVs and 818,401 common indels detected using GATK, to allow for a fair comparison of the contribution of different variant types. We used FastQTL v2.184 (Ongen et al. 2016) to perform *cis*-eQTL mapping, customized to accomidate the unique architecture of SVs (Chiang et al. 2017), using a *cis* window of 1 Mb on either side of the TSSs of autosomal and X-chromosome genes with a permutation analysis to identify the most significant marker for each gene. For each tissue we applied the same covariates described in Chiang et al. 2017. We corrected for multiple-testing at the gene-level using the Benjamini-Hochberg method with a 10% FDR.

### Feature enrichment

To evaluate whether SVs that cause common gene expression changes are enriched in particular genomic features, we calculated a previously described causality score (Chiang et al. 2017) generated by taking the product of the SV heritability fraction obtained from GCTA (Yang et al. 2011) and the causal probability generated by CAVIAR (Hormozdiari et al. 2016) for the strongest-associated SV within the *cis* region of each eQTL. No associated SVs were identified in 199 eQTLs due to the subset of samples with available data in the relavent tissue and thus were not included in enrichment analyses. GCTA heritability estimates could not be calculated for a small number of eQTLs (6,146/299,187) due to nonpositive definite matrices, likely resulting from small sample sizes, and these loci were excluded from feature enrichment analyses. For SVs that were associated with multiple eQTLs or the same eQTL in multiple tissues, we selected the eQTL (tissue/gene pair) for which the SV had the highest causality score. SVs were allocated into bins based on causality score quantiles, with the first bin consisting of SVs in the bottom 50% of causality scores and the other five consisting of deciles of the top 50% of scores.

Next, we counted the number of SVs in each bin that intersected with various genomic annotations. We allowed 1 kb of flanking distance surrounding all annotations with the following exceptions: GENCODE exons, no flanking distance; proximity to TSS and 3’ gene end, 10 kb of directional flanking distance; topologically associated domain boundaries, 5 kb of flaking distance; Roadmap Epigenomics segmentation states, no flanking distance. SVs associated with multiple eGenes were considered to touch an eGene if they overlapped with the exons of any associated gene. SVs that touched an exon of an associated eGene were excluded from all feature enrichment analyses except for the enrichment of affected eGenes. To generate a shuffled null for comparison, SVs within each causality bin were shuffled with BEDTools v2.23.0 (Quinlan and Hall 2010) into non-gapped regions of the genome within 1 Mb of the TSS of a gene. We calculated the fold enrichment of the number of SVs that intersect with each genomic feature compared to the median number of intersections observed for 100 randomly shuffled sets within each causality bin. These shuffled sets were also used to empirically derive the 95% confidence intervals.

Regions 10 kb upstream of TSS and downstream of 3’ gene end were defined based on GENCODE v19 gene positions. DNAse hypersensitive regions and enhancer regions with a minimum support of 2 were obtained from the Dragon ENhancers database (DENdb) (Ashoor et al. 2015). We downloaded FunSeq 2.1.0 (Fu et al. 2014) regions and topologically associated domain boundaries from human embryonic stem cells from author websites (http://archive.gersteinlab.org/funseq2.1.0_data/ and http://compbio.med.harvard.edu/modencode/webpage/hic/hESC_domains_hg19.bed). GeneHancer (Fishilevich et al. 2017) enhancer regions for b38 were downloaded from the UCSC genome browser (Kent et al. 2002) and lifted over to b37 using CrossMap v0.2.6 (Zhao et al. 2014). Regions defined by the ENCODE (Encode Project Consortium 2012) project were downloaded from the UCSC genome browser. To evaluate the intersection with the chromatin segmentation state annotations from the Roadmap Epigenomics Project (Kundaje et al. 2015), we downloaded the core 15-state model annotations for all 127 available epigenomes (https://egg2.wustl.edu/roadmap/data/byFileType/chromhmmSegmentations/ChmmModels/coreMarks/jointModel/final). We used BEDTools multiIntersectBed (Quinlan and Hall 2010) to identify genomic intervals where each of the 15 annotations is found in at least 10 of the 127 available epigenomes and used these collapsed regions as the annotation intervals for SV intersections.

### eQTL tissue specificity

We selected significant gene-variant pairs identified in eQTL mapping with available expression data available across all 48 tissues in which eQTL analyses were performed. These pairs were only required to have a significant eQTL in one tissue. We used METASOFT v2.0.0 (Han and Eskin 2011) to perform a meta-analysis of the selected eQTL effect sizes and their standard errors across all 48 tissues. METASOFT employs a mixed effects model (RE2) to generate a posterior probability that an effect exists in each tissue (*m*-value) (Han and Eskin 2012). To allow computational feasibility with the relatively large number of tissues sampled, the Markov Chain Monte Carlo (MCMC) method was used to approximate these values. The *m*-values generated indicate whether a tested eQTL is active (*m*>0.9), inactive (*m*<0.1), or has ambiguous activity (0.1≤*m*≤0.9). Only eQTLs with at least 43 tissues having known (active or inactive) activity were included in analyses. eQTLs with active status in at least 75% of tissues with known activity were defined as “constitutively active.”

### Identification of expression outliers

We performed Z transformation of PEER-corrected expression values without quantile normalization across the 53 tissues for which RNA-seq data was available from the GTEx consortium. We limited analysis to the 513 European individuals, the largest subpopulation in the cohort, who had available WGS data. We defined two sets of gene expression outliers (gene/sample pairs) among these individuals: “multi-tissue” expression outliers in which an individual’s absolute median Z-score of a gene’s expression across all available tissues was ≥2, as previously described in (Chiang et al. 2017), and “tissue-restricted” outliers in which an individual’s absolute Z score for a gene’s expression was ≥4 in at least two different tissues. The two tissue requirement was necessary to eliminate false positive expression outliers resulting from individual tissues with systematically aberrant gene expression profiles for an individual. Additionally, we defined a set of control gene/sample pairs in which an individual’s absolute Z score of a gene’s expression was less than 1 across all tissues for which RNA-seq data was available. For all definitions we limited to gene/sample pairs with data available in at least 5 tissues. We removed one individual (GTEX-14753) from this analysis due an excessive number of expression outliers.

### Rare variant association with expression outliers

We identified 15,016 structural variants that were positively genotyped in no more than one individual in the European cohort. Because large rare structural variants tend to affect gene expression through dosage changes, we removed 12 variants larger than 1 Mb in size from this analysis. We calculated the enrichment of singleton SVs overlapping with multi-tissue outlier transcripts and the flanking 5 kb sequence by randomly shuffling the outlier individual names 1,000 times to determine the median number of times a rare variant randomly co-occurred with an outlier, as described in (Chiang et al. 2017). We also performed the reciprocal analysis counting the number of outliers that co-occurred within 5 kb of a rare SV. We repeated these calculations for increased outlier-flanking regions of 10 kb, 25 kb, 50 kb and 100 kb.

### Feature enrichment for outlier-associated SVs

We performed intersections between the 369 noncoding outlier-associated SVs and the same genomic features and chromatin segmentation states evaluated for eSVs. The above intersections were repeated for the 1,416 noncoding control-associated SVs and we calculated the fold enrichment of outlier-associated SVs in each feature compared to control-associated SVs.

### Regional effect of rare SVs

To evaluate the broader regional effects of rare, gene expression-altering SVs, we calculated the number of tissue-restricted outlier genes, referred to as “primary” outliers, located in the spanning region and 1 Mb of flanking sequence both upstream and downstream of the 469 SVs previously identified as being associated with an expression outlier. We repeated this analysis with a relaxed definition of tissue-restricted expression outliers, referred to as “secondary” outliers, in which the absolute Z score cutoff was reduced from |Z|≥4 to |Z|≥3. We compared the number of primary and secondary outliers found in the expanded region surrounding outlier-associated SVs to the expanded region surrounding the 1,224 control-associated SVs. Finally, because the controls defined above do not represent a null expectation, we performed 1,000 random permutations of the outlier-associated SV sample names and calculated the median number of associated primary and secondary outliers for each SV in order to determine how frequently rare expression-altering SVs co-occurred with primary and secondary outliers in random individuals.

### DATA ACCESS

The SV call set is available under dbGaP accession code [TBD].

## ACKNOWLEDGMENTS

The authors thank E.J. Gardner for advice on MELT and R.E. Handsaker for advice on Genome STRiP. This work was supported by a Mr. and Mrs. Spencer T. Olin Fellowship for Women in Graduate Study (A.J.S.) and by the NIH/NHGRI UM1 HG008853 (I.M.H).

